# Fish spawning events stimulate trophic hotspots across freshwater food webs

**DOI:** 10.1101/2025.09.30.679526

**Authors:** Timothy J. Fernandes, Kayla R. S. Hale, Brett M. Studden, Royce Steeves, Sofia Pereira, Jonathan B. Armstrong, Kevin S. McCann, Trevor Middel, Martyn E. Obbard, Derek Potter, Mark Ridgway, Brian J. Shuter, Tyler D. Tunney, Bailey C. McMeans

## Abstract

Mass reproductive events, where large numbers of organisms aggregate for synchronous reproduction, are often captivating and ecologically significant, boosting offspring survival and reducing time spent searching for mates. However, mass reproduction can also instigate cryptic but consequential responses across entire food webs. Reproductive materials are abundant and accessible resources that can attract mobile consumers from up to thousands of kilometers away. Yet, their consumption, especially in aquatic systems, is difficult to detect and rarely characterized. Here, we combine molecular techniques with acoustic telemetry, literature review, and extensive natural history observations to investigate the food web consequences of synchronized reproduction in freshwater fishes. First, we demonstrate that a common but underappreciated fish species, white sucker, creates a resource pulse used ubiquitously by consumers, from local invertebrates and fishes to mobile predatory fishes, birds, and terrestrial mammals. Spawning white sucker create trophic hotspots that attract consumers across trophic levels and ecosystems to feed on eggs, spawning adults, and aggregated egg predators. Then we show that egg provisioning and predation is widespread among north-temperate freshwater fish species, highlighting that resource pulses instigated by mass reproduction may play a critical but underappreciated role in freshwater and terrestrial ecosystems.

## 2.0 Introduction

Synchronized mass-reproductive behaviour is widespread across the Tree of Life. Across taxa, synchronized reproductive behaviour can reduce the time spent mate searching and also improve offspring survival, as mass gamete deposition swamps potential predators (Ims, 1990b, 1990a; Janzen, 1967). Yet, the general role of these events as critical resource pulses that shape the spatiotemporal dynamics of trophic interactions and energy fluxes across ecosystems is poorly described. While prey are generally invulnerable or inaccessible, reproductive events offer rare opportunities for predators to exploit vulnerable and accessible prey resources (e.g., spawning adults and their gametes) that become concentrated in space and time (Abrams & Walters, 1996; Magnhagen, 1991). The potential magnitude of such reproductive resource pulses has been well-documented, as exemplified by strikingly-coloured Pacific salmon spawning runs in coastal tributaries (Armstrong et al., 2016; Gende et al., 2002; Schindler et al., 2003) and synchronized masting events that flood the forest floor with nuts and other seeds (Beck et al., 2024; Ostfeld & Keesing, 2000; Selva et al., 2012). Examples of reproductive events that offer unique, spatiotemporally concentrated opportunities for resource acquisition abound in ecology, spanning ecosystems and taxa (Deakos et al., 2011; Fuiman et al., 2015; Gilbert et al., 2025; Rayl et al., 2015). These high-resource events are likely central to attracting and supporting diverse consumer communities (Abrahms et al., 2021; Beck et al., 2024; Yang et al., 2008), shaping the functioning of food webs and driving general patterns in consumer behaviour, physiology, and abundance (Bailey et al., 2019; Naiman et al., 2002; Ruff et al., 2011). Nonetheless, we have a surprisingly limited understanding of how these events may attract species from local to regional scales, across traditional habitat boundaries to impact whole food webs that ignore spatial boundaries (e.g., freshwater and terrestrial).

In even the largest freshwater systems globally, the Laurentian Great Lakes, the reproductive behavior of native fishes can structure the dynamics of local primary production (Childress et al., 2014) and the annual energy budgets of mobile predators (Stockwell et al., 2014). The annual spawning migrations of inland freshwater fishes can promote extensive energy and nutrient transport across ecosystem boundaries, likely representing critical fluxes for some of the most at-risk food webs globally (Dudgeon et al., 2006; Reid et al., 2019; Tickner et al., 2020). For example, the spring migration of white sucker (*Catostomus commersonii*) from productive lake habitats into oligotrophic spawning streams can increase the primary and secondary productivity of recipient streams multiple fold by depositing carbon and other limiting nutrients through reproductive material and excrement (Childress & McIntyre, 2015). Catostomids, including white and longnose sucker, are among the most widespread migratory fishes in North America, commonly representing the largest proportion of fish biomass in its freshwaters (Becker, 1983). However, suckers are non-game species that are generally thought to have low recreational and commercial value, such that their contributions to freshwater food web dynamics are poorly documented across much of their range (Cooke et al., 2005; Murchie et al., 2024). Because of their abundance, broad geographic distribution, and extensive seasonal spawning aggregations, suckers represent a valuable set of focal species for understanding the extent to which fish spawning can concentrate trophic interactions and mediate consumer behaviour via hotspots of critical energy and nutrient flow (i.e., trophic hotspots). Indeed, spawning in other freshwater fishes may also have the capacity to provision similar trophic hotspots through reproductive aggregations and nutrient deposition, though this capacity is currently poorly described.

Across freshwater fish species, the development of eggs and reproductive material represents one of the most energetically costly life history processes of any given year (Ginther et al., 2024). To improve the probability of offspring survival, developing eggs are stocked with a cocktail of limiting macromolecules including essential poly- and mono-unsaturated fatty acids, lipoproteins, and amino acid residues that then fuel and facilitate larval development (Migaud et al., 2010; Reading et al., 2018; Wiegand, 1996; Wiegand et al., 2014). As a result, eggs are a high-quality food source with energy densities 2-3-times greater than even alternative high-quality prey (e.g., predatory macroinvertebrates; Stockwell et al., 2014). Further, many large- and small-bodied freshwater fishes exhibit high fecundity (e.g., >5 000 eggs per kg in mature females; Vélez-Espino et al., 2006; Winemiller & Rose, 1992), such that vast numbers of eggs are deposited during spawning events to swamp potential egg predators (i.e., ‘egg boons’; Fuiman et al., 2015; Ims, 1990a). Unless specifically defended as seen in some salmon (Armstrong et al., 2010), eggs are easily handled by predators due to their small size (alleviating gape-limitations in even small-bodied predators; Fox, 1978; Stauffer & Wagner, 1979), high nutritional quality (reducing digestion times; Hunter et al., 2012), predictability (synchronously cued by environmental conditions; Ims, 1990a), and density at spawning locations (reducing search and attack times; Richardson et al., 2011). This allows egg predators to gorge on eggs, exceeding consumption levels observed even during other resource pulses they encounter (e.g., emergence of adult invertebrates; Richardson et al., 2011; Wesner, 2016). In this way, energy initially integrated into and bound by individual consumers may be efficiently redistributed across trophic levels of the entire food web via egg predation, for example when eggs of top predators (e.g., lake trout *Salvelinus namaycush*) are consumed by low-level consumers (e.g., white sucker *Catostomus commersonii*) in a “counter gradient” flow of energy (Fuiman et al., 2015; Wasylenko et al., 2013).

However, detecting and quantifying the consumption of reproductive material can present many logistical challenges that limit our ability to describe its generality in nature. For many consumers, fish eggs and reproductive material are available for only a short window of time, so spatiotemporally informed and targeted monitoring is required to detect egg predation events. In many cases lacking indigestible material, fish eggs themselves degrade rapidly in the digestive tracts of consumers, limiting opportunities for detection (Caroffino et al., 2010). For example, newly hatched smelt embryos are considered unrecognizable via conventional stomach content analyses within two hours of consumption at 11°C, with eggs expected to be digested even more rapidly (Crowder, 1980). Additionally, much of the existing work documenting egg predation by fishes has focused on documenting early life stage mortality of culturally-important fishes through conventional stomach content analyses (Lutz et al., 2020; Roseman et al., 2006; Schneider, 1997). This approach likely underestimates the extent of egg predation as well as the diversity of egg predators present. Identifying egg predation by invertebrate consumers is similarly challenging, commonly relying on physical observations in the lab or inferring consumption from unprotected benthic egg baskets or stable isotope signatures (Caroffino et al., 2010; Childress et al., 2014; Fox, 1978). As such, the extent to which consumers across trophic levels and ecosystem boundaries exploit reproductive resource pulses generated by freshwater fishes remains largely unknown.

Here, we used a combination of approaches to investigate (1) if fish spawning aggregations represent trophic hotspots characterized by a reproductive resource pulse to diverse consumers in recipient habitats and behavioural responses by mobile consumers; and (2) if spawning aggregations are likely to provision reproductive resource pulses to north-temperate freshwater fishes more broadly. First, we conducted a field study in a north-temperate lake food web (Smoke Lake, Ontario, Canada) to characterize the exploitation of a reproductive resource pulse generated by one of the most widespread migratory freshwater fishes in North America, the white sucker. Using both standard stomach content analyses and molecular techniques, we documented the consumption of reproductive material during sucker spawning in nine species of freshwater fish and six orders of benthic macroinvertebrates. Simultaneously, we used acoustic telemetry to study the behavioral response of a mobile freshwater predator in Smoke Lake (smallmouth bass, *Micropterus dolomieu*), showing that even under stressful thermal conditions, this predator tracked the reproductive resource pulse in space and time. We additionally synthesized eight years of standardized natural history observations from a nearby lake system, finding evidence of an extensive community of terrestrial and aquatic species consuming spawning white sucker adults and reproductive material. Finally, we conducted a literature synthesis to quantify the extent to which north-temperate fish species can be classified as predators or provisioners of reproductive resources. By combining these data sources, we demonstrate that there is broad potential for the exploitation of reproductive resource pulses introduced by spawning aggregations of freshwater fish across multiple trophic levels in freshwater and terrestrial food webs. We also find that egg predation is a widespread phenomenon among north-temperate fishes. We conclude with a discussion in which we assert that reproductive resource pulses provisioned by spawning fish aggregations attract consumers and concentrate energy flows in space and time – a trophic hotspot shaping freshwater and terrestrial food webs.

## 3.0 Results

### 3.1 Overview of field study – characterizing the consumer community in Smoke Lake, Ontario

To characterize the potential for fish eggs and reproductive material to represent an impactful resource pulse for consumers across trophic levels, we conducted an observational field study targeting primary consumers to top predators in Smoke Lake, Algonquin Provincial Park, Ontario, Canada. Smoke Lake hosts a common cold-water, north-temperate fish community comprised of lake trout, brook trout (*Salvelinus fontinalis*), cisco (*Coregonus artedi*), smallmouth bass (*Micropterus dolomieu*), burbot (*Lota lota*), lake whitefish (*Coregonus clupeaformis*), yellow perch (*Perca flavescens*), white sucker, golden shiner (*Notemegonus crysoleucas*), common shiner (*Luxilus cornutus*), creek chub (*Semotilus atromaculatus*), blackchin shiner (*Notropis heterodon*), blacknose shiner (*Notropis heterolepis*), bluntnose minnow (*Pimephales notatus*), brown bullhead (*Ameiurus nebulosus*), northern pearl dace (*Margariscus margarita*), northern redbelly dace (*Chrosomus eos*), and finescale dace (*Chrosomus neogaeus*); white sucker have been observed to migrate and spawn in the creek deltas of Smoke Lake.

We selected three sampling sites located at the mouths of tributary streams entering the lake: Site 1, a negative control site where no aggregations of adult white suckers or eggs have been observed, Site 2, where migrating white suckers have been observed but access to upstream spawning habitat is blocked by a large beaver dam such that neither spawning behaviour nor egg deposition has not been observed, and Site 3, where both migrating white suckers and eggs have been observed. As expected from previous observations, we observed no adult white sucker at Site 1, adult white sucker aggregations but no spawning behavior at Site 2, and spawning white sucker aggregations with visual confirmation of egg deposition at Site 3 (Figure 1). We also used telemetry data from an existing acoustic array in Smoke Lake to detect daily movements of tagged smallmouth bass (n = 19 unique individuals) and white sucker (n = 15 unique individuals) at each site prior to, during, and following sucker spawning.

**Figure 1.**
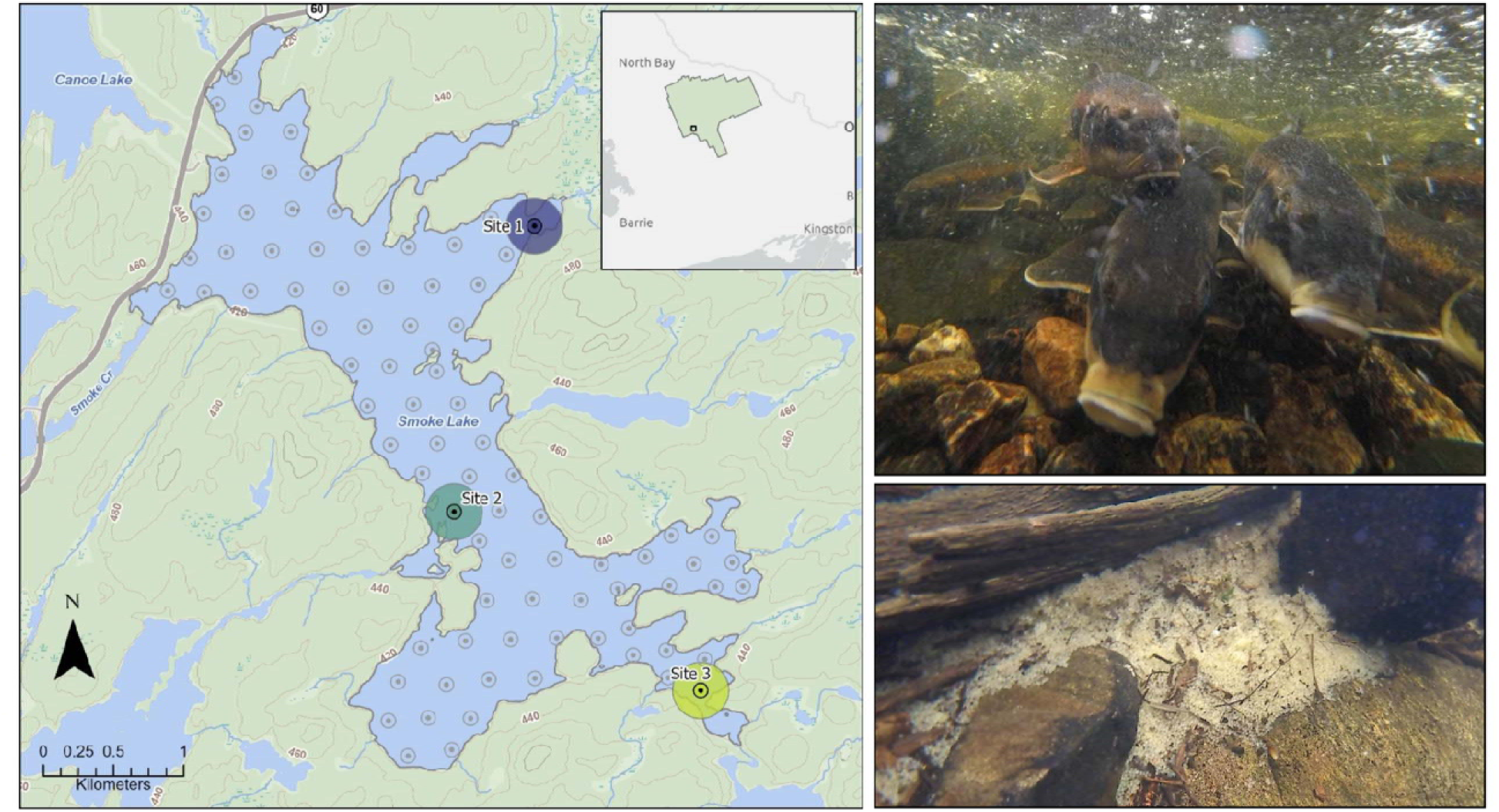
Field sites with and without white sucker spawning activity. Map of Smoke Lake, Ontario, Canada (left) depicting the array of acoustic telemetry receivers (gray dots) and field sampling sites (colored circles). The detection range of each receiver is approximately equal to the radius of the colored circles (for clarity, only the ranges of sampling-site specific receivers are shown). Purple (top): Site 1, no white suckers observed; Teal (middle): Site 2, white sucker spawning aggregation confirmed, but no eggs observed; Green (bottom): Site 3, white sucker spawning aggregation confirmed (top right) with eggs observed throughout the lakebed (bottom right).

At each sampling site, we collected fish and invertebrates for approximately one week after spawning aggregations of white suckers were visible. We sampled all fish species captured across all sites (n = 9 species: bluntnose minnow, brown bullhead, burbot, common shiner, creek chub, northern pearl dace, northern redbelly dace, smallmouth bass, yellow perch). We focused our macroinvertebrate collections on Trichoptera, Plecoptera, Odonata, Ephemeroptera, Megaloptera, and Decapoda, where available. The stomach contents of collected fish were identified to the finest possible taxonomic resolution following gastrointestinal dissection; for smallmouth bass, we supplemented dissection data with stomach contents gathered via gastric lavage of non-lethally sampled individuals. For both fish and invertebrates, we then used quantitative PCR (qPCR) to amplify and detect white sucker DNA in gastrointestinal contents extracted via aseptic dissection.

### 3.2 Feeding on reproductive resources is cryptic but present across consumers and trophic levels

We failed to detect egg predation in Smoke Lake using traditional methods of stomach content analysis including gastric lavage (n = 27) of large-bodied fish (smallmouth bass) and gastrointestinal dissection (n = 171) of small- and large-bodied fishes across all sites. However, by using qPCR, we were able to detect white sucker DNA in the stomach and intestinal contents in all the fish species (n = 8) and the macroinvertebrate orders (n = 5) collected at sites where sucker eggs (egg diameter: 3.0-3.3 mm, Fuiman & Trojnar, 1980; Manny et al., 2010; egg dry mass: 1.4 mg/egg; Johnston, 1997) were confirmed visually, except in ephemeropterans (Site 3; Figure 2). At Site 3, 70% of all fish sampled contained detectable levels of sucker DNA in their digestive tracts (n = 72 of 101), compared to 19% at Site 2 (n = 11 of 58) and 8% at Site 1 (n = 1 of 12; Figure 2). At Site 3, 68% of all macroinvertebrates sampled contained detectable levels of sucker DNA (n = 31 of 46), compared to 45% at Site 2 (n = 5 of 11) and 2% at Site 1 (n = 1 of 39; Figure 2A). Though the consumption of white sucker carcasses could also have contributed to observed DNA signals, our sampling occurred during the beginning of spawning at which time no carcasses were detectable during both snorkelling and boat-side observations. Further, passive ingestion of detectable levels of white sucker DNA appears unlikely given the absence of signals (i.e., individuals with no detectable DNA in their digestive tracts) in all invertebrate orders and all fish species with more than two individuals sampled.

**Figure 2:**
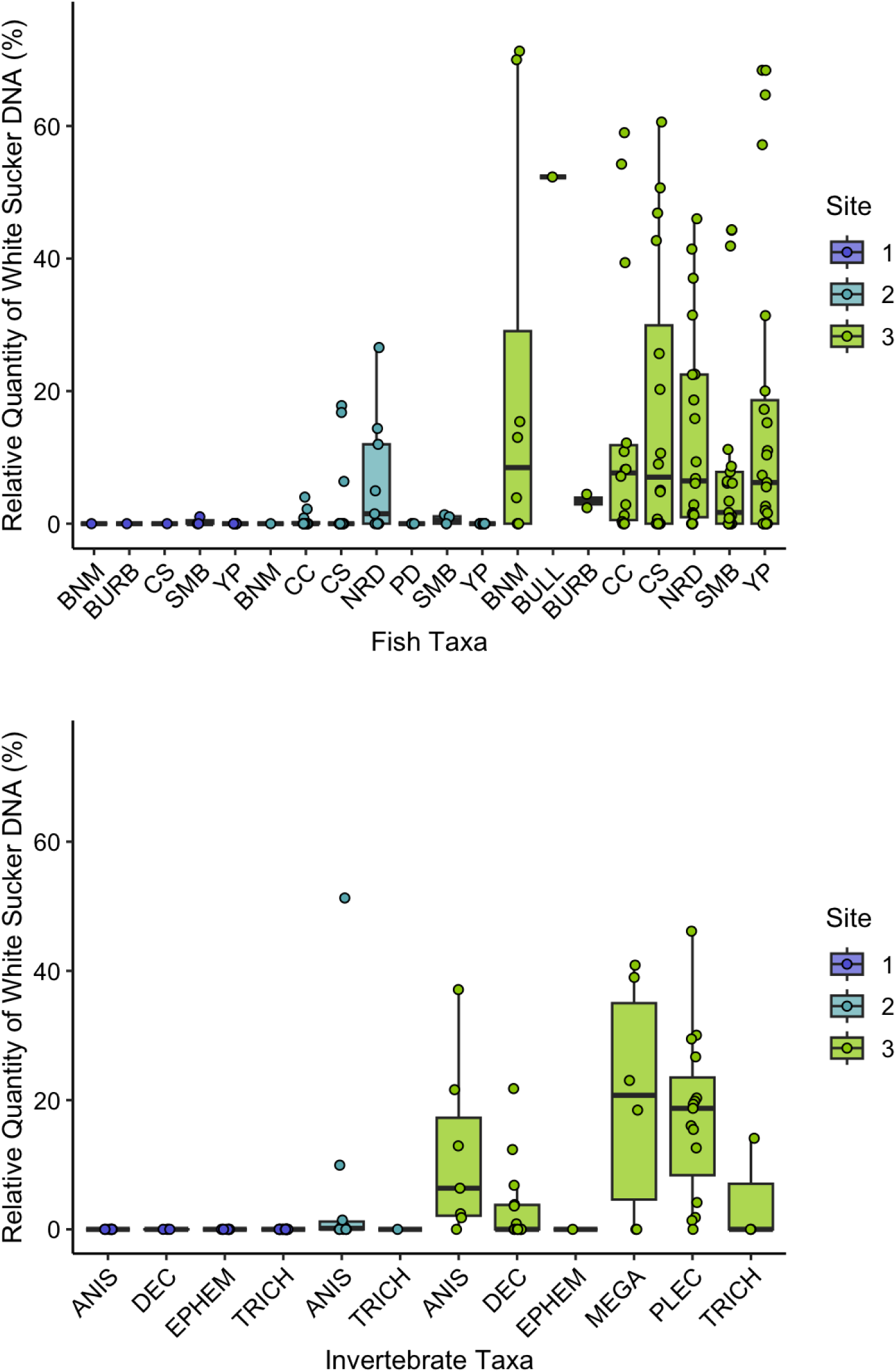
Diverse fish and macroinvertebrate taxa predate white sucker eggs. Relative quantity of white sucker DNA detected in the digestive tract of macroinvertebrates (top; ANIS - Anisoptera, DEC - Decapoda, EPH - Ephemeroptera, MEGA - Megaloptera, PLEC - Plecoptera, TRI - Trichoptera) and fish (bottom; BNM - bluntnose minnow, BULL - brown bullhead, BURB - burbot, CC - creek chub, CS - common shiner, NRD - northern redbelly dace, PD - pearl dace, SMB - smallmouth bass, YP - yellow perch) across three sites in Smoke Lake, Algonquin Park, Ontario. Box plots show median and quantiles of observations, colored by sample site. Purple (left): Site 1, no suckers observed; Teal (middle): Site 2, sucker aggregation confirmed; Green (right): Site 3, sucker aggregation and eggs confirmed.

Among macroinvertebrates at Site 3, individuals of Plecoptera, Anisoptera, and Megaloptera were the most likely to be egg predators as informed by qPCR, though Decapoda and Trichoptera also demonstrated some degree of egg consumption (Figure 2). We detected similar quantities of white sucker DNA in all fish species analyzed at Site 3, though smallmouth bass also consumed macroinvertebrate species that could have contributed white sucker DNA indirectly. No juvenile suckers were detected at any of the sites or in any bass stomachs, suggesting that the detected sucker DNA was due to consumption of eggs or consumption of egg-feeding invertebrates. When sucker aggregations were absent (Site 1), white sucker DNA was not detectable in any of the sampled consumers except smallmouth bass, a highly mobile predator. However, when sucker aggregations were present but prevented from accessing upstream spawning habitat by a natural barrier (beaver dam; Site 2), Anisoptera and several species of prey fishes (common shiner, creek chub, northern redbelly dace) were still able to exploit sucker eggs. Here, spawn-ready suckers that aggregated beneath the beaver dam likely released small numbers of eggs that were not detected by observers but remained available to some of the local consumer community. Thus, the reproductive resource pulse provisioned by white sucker in Smoke Lake was exploited by nearly the entire consumer community (12 out of 13 fish and invertebrate taxa) and by over 70% of both fish and invertebrate consumers (103 out of 147 sampled individuals) present at sites where the pulse was available.

### 3.3 Mobile top predators track reproductive resource pulses

Analysis of the telemetry data revealed that smallmouth bass tracked the sucker spawn in both space and time. Detections of individually tagged smallmouth bass and white sucker at the Site 3 receiver were patterned synchronously from late winter to early summer (lag = 0, r = 0.74, P < 0.05). This window of time captured the arrival, residence, and departure of both white sucker and smallmouth bass. Though smallmouth bass are not generally expected to be active below 10°C (Munther, 1970; Shuter et al., 1980), individuals began entering the site when lake temperatures remained below 5°C (Figure 3). In total, nearly 80% of the tagged smallmouth bass population (n = 14 of 19) were detected at the receiver nearest to Site 3 during the sucker migration and spawn. Using telemetry receivers present in surrounding lakes, we were also able to identify one individual that travelled through two connected lakes (Canoe Lake, Tea Lake) and across the entire length of Smoke Lake before being detected at Site 3, traversing a minimum distance of 10 km of lake and creek reaches in less than 48 hours.

**Figure 3.**
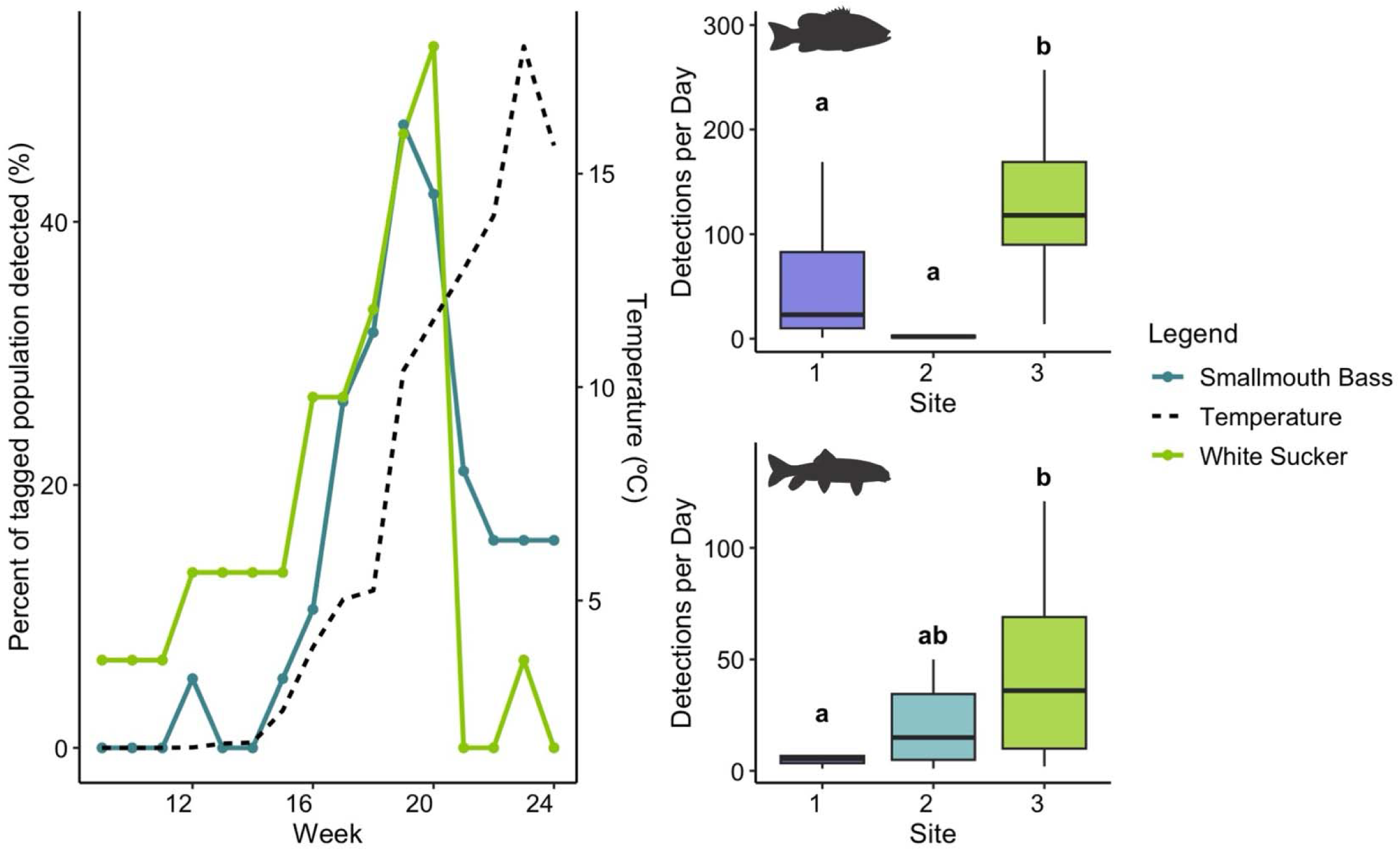
Smallmouth bass track white sucker spawning aggregations in space and time. (A) The percentage of all tagged smallmouth bass (n = 19) and white sucker (n = 15) in Smoke Lake detected weekly between March and July 2023 at the single acoustic telemetry receiver located closest to the Site 3 creek inflow. The dashed lined illustrates the mean weekly temperature as measured by an Onset HOBO Pendant temperature logger co-located with the Site 3 receiver. (B) Detections of tagged smallmouth bass (top) and white sucker (bottom) across three sites in Smoke Lake. Purple (left): Site 1, no suckers observed; Teal (middle): Site 2, sucker aggregation confirmed; Green (right): Site 3, sucker aggregation and eggs confirmed. Shared lettering indicates similarity in the number of detections between sites as informed by post-hoc Dunn’s multiple comparisons test using a Bonferroni p-value adjustment.

Across the three sites, white sucker detections agreed with observations of sucker aggregations. Detections were greatest at Site 3, followed by Site 2, with no detections recorded at Site 1 (Kruskal-Wallis ANOVA; H(2) = 9.69, p = 0.008). Similarly, smallmouth bass detections were greatest at Site 3, followed by Site 1, with no detections recorded at Site 2 (H(2) = 33.38, p < 0.0001). Thus, the local hotspot of food web activity provisioned by white sucker also attracted mobile predators, like smallmouth bass.

### 3.4 Both terrestrial and freshwater consumers exploit sucker spawning migrations

To quantify exploitation of white sucker reproductive resource pulses by terrestrial consumers, we compiled 8 years of standardized natural history observations targeted during sucker spawning in Wright Creek where it feeds into Opeongo Lake, Algonquin Provincial Park, Ontario, Canada (∼30 km from Smoke Lake). From 2008-2015, observations were collected from an observation blind and a network of camera traps upstream of the mouth of Wright Creek during sucker spawning season, yielding 3343 wildlife observation records from images and 687 wildlife observation records from primary observation. From these, we identified 17 bird and 5 mammal species repeatedly visiting and foraging at the site of sucker spawning over the 8-year period (Figure 4). Waterbirds and piscivorous birds were the most commonly observed bird species, with common mergansers and herring gulls together accounting for over 50% of blind and camera trap observations and both identified across all years (Supporting Information, Table S1, Table S2). Common ravens (5.6-8% of observations), bald eagles (3.6-7% of observations), and dabbling ducks (1-12.8% of observations) were also common and consistently observed across years. Then, raccoons (*Procyon lotor*), American black bears (*Ursus americanus*), white-tailed deer (*Odocioleus virginianus*), and mustelids were the most commonly observed mammals (Supporting Information, Table S1, Table S2). When present, direct egg consumption was confirmed by observer notes.

**Figure 4.**
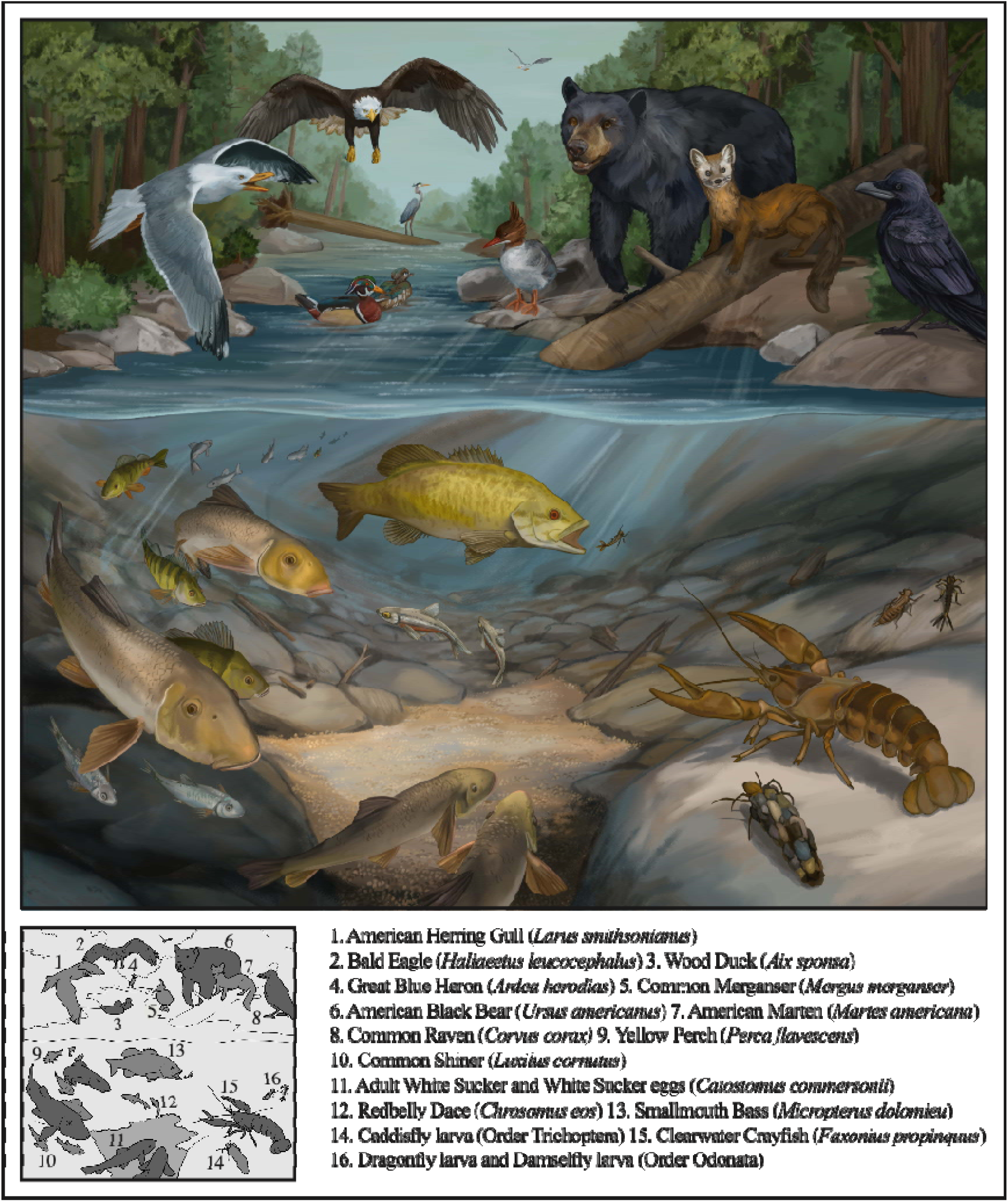
The extensive consumer community attracted by white sucker spawning. An illustration of the extensive consumer community that appears to aggregate around white sucker spawning locations, with observations compiled from both Smoke Lake and Opeongo Lake, Algonquin Provincial Park, Ontario, Canada. Only a subset of species that were observed to exploit the sucker spawning run are pictured here. Artwork by Sophia Bos.

### 3.5 Exploitation of spawning migrations is common in north-temperate freshwater fish

To assess the extent that reproductive resource pulses potentially contribute to local freshwater food webs, we conducted a literature review and synthesis focusing on documented egg predation among northern freshwater fishes (search string in Google Scholar and Clarivate Web of Science: “(“egg predator” OR “egg predation”) AND freshwater AND fish”). From the literature reviewed (see Methods), we included only the n = 42 studies with direct evidence for egg predation events (i.e., both predator and prey species identified) in wild fishes from north-temperate (> 30°N) freshwater systems.

These studies document egg predation by 43 species of north-temperate freshwater fish from 16 taxonomic families (Supporting Information, Table S3), with body sizes spanning four orders of magnitude. Of the 16 families identified, catostomids and centrarchids fed on the most diverse set of prey species (Figure 5). Fish species that provide egg resources (i.e., egg provisioners) were also highly diverse, with 34 freshwater fish species across 12 families provisioning eggs to consumers. Of these, catostomids (n = 6 species), salmonids (n = 12 species), and centrarchids (n = 5 species) represented the majority (68%) of documented egg-provisioning species. Interestingly, spawning and embryo development time varied widely across egg provisioning species (Figure 5B). Spring spawning prey species exhibit substantially shorter embryo development periods (2-38 days) than fall and winter spawners (92-140 days), suggesting that the temporal window of egg availability differs substantially among prey species. Thus, the diversity of egg-predator and egg-provisioner fishes results in temporal heterogeneity in the availability of reproductive resources.

**Figure 5:**
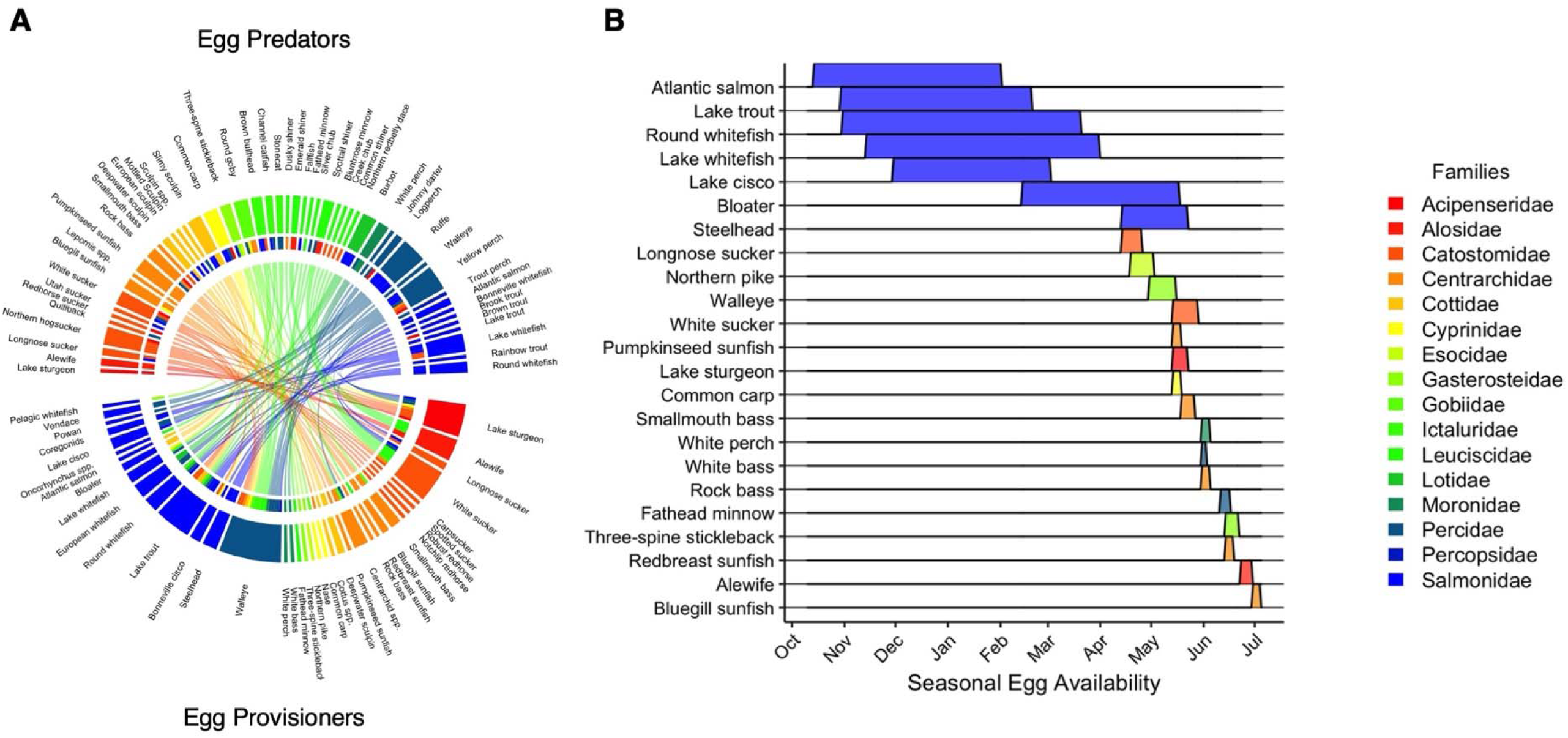
Egg predation is a generalized strategy among north-temperate fish species, creating a resource portfolio for mobile consumers. (A) Chord diagram summarizing literature review of egg predation records among north-temperate fishes. Egg predators (top) are linked to egg provisioners (bottom). (B) Range plot showing potential seasonal availability of eggs provisioned by fish species from our literature review that co-occur in North American freshwater systems. Colors correspond to taxonomic family, following. Ranges show the date of expected spawning onset and span the expected length of embryo development to emergence extracted from Scott and Crossman (1998). These dates vary across latitude and populations; here, we plotted midpoints of dates reported in Scott and Crossman (1998) simply to illustrate the approximate timing and length of resource availability to egg predators.

## 4.0 Discussion

Here, we demonstrate that freshwater fishes provision reproductive resource pulses that are exploited by consumers at multiple trophic levels of both aquatic and terrestrial food webs, from primary consumers to top predators. We widely sampled the local consumer community of an inland lake using quantitative PCR (qPCR) to amplify white sucker DNA from consumer digestive tracts. When eggs were available prey items, all sampled taxa (eight fish species and six macroinvertebrate orders) contained detectable levels of sucker DNA except Ephemeroptera. Additionally, we used telemetry to document the movements of white sucker and a top predator fish, smallmouth bass (SMB). SMB appeared to track the availability of sucker eggs in both space and time, even under seemingly prohibitive thermal conditions (i.e., between 5 and 10°C prior to and during egg availability, temperatures thought to limit activity and feeding). Finally, we presented eight years of local natural history records demonstrating direct and repeated consumption of aggregated adult white sucker or their reproductive material by 17 bird and five terrestrial mammal species. Together, our evidence suggests that white sucker spawning aggregations create a trophic hotspot in freshwater and adjacent terrestrial habitats, characterized by a spatially and temporally constrained pulse of resources exploited by consumers across feeding guilds, from herbivores to top predators. Importantly, our review then demonstrates the potential for such hotspots across north-temperate freshwater fishes and ecosystems: in addition to white sucker, spawning behavior by many other fish species also provision resource pulses for diverse fish species not typically regarded as egg predators. The ubiquity of synchronized spawning events across inland freshwaters suggests that freshwater fishes are likely strong mediators of trophic hotspots across entire watersheds.

In what follows, we first discuss how fish eggs provide critical but underappreciated pulsed resources that subsidize freshwater food webs, with effects that cascade across the landscape. Then, we compare this phenomenon to better studied examples in marine systems, resource waves more broadly, and even masting events by long-lived plant species in terrestrial ecosystems. We contend that the spatiotemporal heterogeneity of egg deposition across coexisting fish species staggers reproductive resource pulses in space and time, likely cascading through local food webs to create a ‘portfolio’ of reproductive resource pulses at the landscape scale.

### 4.1 Reproductive resources are exploited by species across diverse trophic guilds

Our field study demonstrated that white sucker reproductive resources provided a pulsed energy source for a range of local basal consumers and mobile top predators. Though large, well-defended fish eggs (e.g., salmonid eggs) can be detected in fish stomachs hundreds of metres away from sites of egg deposition (Armstrong et al., 2013), the eggs of many fish species are rarely detectable soon after consumption (Lutz et al., 2020) by conventional methods (gastric lavage, gastrointestinal dissection). By widely sampling with qPCR, we provided novel evidence of direct consumption of freely available reproductive resources by diverse benthic macroinvertebrate consumers, from caddisfly and dragonfly larvae to dobsonfly larvae and crayfish (Caroffino et al., 2010; Fox, 1978). Similarly, our identification of sucker egg consumption by diverse fish species (e.g., an offshore benthic predator burbot, a nearshore predator smallmouth bass, small-bodied nearshore prey fishes) and macroinvertebrates support previous work demonstrating increased growth and abundance of benthic macroinvertebrates and small-bodied fishes in stream reaches that host sucker spawning aggregations (Childress et al., 2014; Jones & Mackereth, 2016). Additionally, our literature synthesis found egg predation to be widespread across freshwater north-temperate fish species. Contrary to the general expectation that small-bodied benthic fishes are the most prevalent egg predators (Fitzsimons et al., 2006; Savino & Henry, 1991), we found evidence of extensive egg predation by fishes across feeding guilds and body sizes. However, targeted sampling of potential egg predators on spawning shoals was only present for a subset of culturally significant north-temperate fish species (e.g., lake trout, lake sturgeon, walleye). Given the difficulty in detecting egg predation events (Lutz et al., 2020), actual numbers of egg predator species likely far exceed our estimate, suggesting that egg predation is a generalized resource acquisition strategy in north-temperate freshwater fishes.

Our synthesis of standardized natural history observations then revealed that many consumers outside of aquatic ecosystems can feed extensively on fish eggs and aggregated spawning adults. For example, across our 8-year observational dataset at Wright Creek, aquatic and terrestrial predators, especially birds (common mergansers, herring gulls, bald eagles, wood ducks, mallards, American black ducks, corvids; Figure 4) and mammals (black bears, raccoons, mustelids; Figure 4), were consistently observed feeding on adult sucker or sucker eggs *in situ*. Though not sampled in our field study, additional aquatic (brook trout, lake whitefish, double-crested cormorants, common mergansers, herring gulls) and terrestrial (common ravens, mustelids) consumers were also repeatedly observed feeding at the site of sucker spawning in Smoke Lake (Site 3; T. Fernandes, *pers. obs.*). Combined with previous evidence collected from a range of taxa (snapping turtles *Chelydra serpentina*: Studden et al., 2025; common mudpuppies *Necturus maculosus*: Harris Jr, 1959), we underscore the sheer diversity of consumers that exploit freshwater fish eggs. The nearly comprehensive consumption of sucker eggs by local and temporarily visiting consumers in our study sites is remarkable, and it is likely that dedicated sampling would reveal a far greater number of terrestrial predators of fish eggs than our localized observations suggest.

Finally, our literature review also demonstrated that 34 freshwater fish species across 12 families also provision eggs to consumers. These species vary in fecundity, nesting behavior, and other defenses associated with reproduction, such that egg accessibility, palatability, and abundance likely mediate the magnitude of the reproductive resource pulse and resultant latency in energy transfer to consumers (i.e., life history-specific trade-offs between predation risk at egg, larval, and juvenile life stages; Childress & McIntyre, 2015). For example, spawning aggregations of white and longnose sucker, species that have relatively small eggs (∼3 mm) and exhibit no nesting or guarding behaviour, provided greater resource subsidies to recipient spawning ground food webs than nest-creating salmonids (Jones & Mackereth, 2016). However, given the low detectability of eggs in digestive tracts, it is likely that recorded provisioning of eggs reflects only widespread and/or high-intensity egg predation that is disproportionately detectable. Stable isotope analyses have proven effective at identifying subsidy assimilation in producers and consumers (Childress et al., 2014; Jones & McKenzie, 2024), though this approach may depend on low-productivity environments where the contributions of other energy channels are reduced. Our work opens new questions about the ubiquity of egg provisioning/predation across freshwater ecosystems, highlighting their potentially critical food web consequences.

### 4.2 Aggregate spawning events concentrate energy flows across freshwater and terrestrial food webs

The deposition of eggs, milt, carcasses, and excrement by migratory freshwater fishes concentrates vast quantities of energy in recipient spawning-ground food webs across freshwater systems (Childress et al., 2014; Jones & McKenzie, 2024; Schindler et al., 2003). In the Laurentian Great Lakes, tributaries that receive resource and nutrient subsidies from migratory fishes, primarily in the form of gametes, excreta, and carcasses, exhibit increased primary and secondary productivity relative to streams and reaches where spawners are absent or excluded (Childress & McIntyre, 2015; Jones & Mackereth, 2016). In marine systems, fish eggs released during mass-reproductive events are thought to contribute critically to essential fatty acid flux through marine food webs (Fuiman et al., 2015), with mobile predators travelling vast distances to aggregate at spawning sites (Hartup et al., 2013; Heyman et al., 2001). We suggest that fish eggs in inland freshwater systems represent similarly important resources that have yet to be appropriately integrated into our understanding of freshwater food webs.

During local spawning events, reproductive fish redistribute energy assimilated through foraging across spatially diffuse habitats to consumers across trophic levels and food web compartments through gamete and waste deposition. We presented data that supports the breadth of this redistribution, with consumption of white sucker eggs observable across the benthic macroinvertebrate community, as well as in terrestrial, avian, and aquatic consumers. This redistribution supports the hypothesis that spawning events drive a “counter-gradient flow” of energy and nutrients, as the eggs of fish become available to all levels of the food web (Fuiman et al., 2015). In addition, the aggregation of consumers across a wide size and trophic spectrum introduces opportunities for efficient foraging by large-bodied predators, feeding on both reproductive material and egg predators themselves. Thus, spawning events instigate trophic hotspots in spatially and temporally explicit pulses that can then have disproportionately strong effects on the annual energy budgets of consumers. For example, cisco eggs can represent 34% of the energy consumed by lake whitefish annually, while being available for less than 20% of the year (Stockwell et al. 2014). The consequences of these flows for food web dynamics (from local to landscape) are understudied but may be both unique and critical for shaping spatiotemporal dynamics in resource quality and species persistence. As these hotspots aggregate lower-level consumers, top predators then have access to both reproductive resources and aggregated egg predators. In cases where top predators preferentially consume aggregated egg predators, the indirect top-down regulation of egg predators may be an adaptive benefit of synchronized reproductive behaviour that requires further study.

### 4.3 Mobile consumers track spawning events as spatially explicit resource pulses

While spatially explicit spawning activity can improve energy flow to and growth of consumers located within spawning habitat patches (Jones & Mackereth, 2016), here we presented evidence that a mobile freshwater consumer (smallmouth bass) tracks sucker egg pulses in space and time. The aggregative response (the rate at which consumers colonize patches with spatially explicit resource pulses; Yang et al., 2008) in tracked smallmouth bass appeared anticipatory, in that fish were detected (visually and telemetrically) at sucker spawning sites prior to observable egg availability, similar to Dolly Varden (*Salvelinus malma*) responsively tracking temporal variation in salmon migrations (Sergeant et al., 2015; Figure 3). Additionally, the relative abundance of bass present at the sucker spawning grounds is noteworthy; at the peak of the bass aggregation in early May, nearly 80% of the tagged smallmouth bass population in Smoke Lake (n = 14/19) was detected at the receiver closest to the sucker spawning run (Site 3). Then, bass appeared to exit the spawning grounds following the sucker outmigration (Figure 3), suggesting that the adult bass population has the capacity to track and exploit the white sucker migration. Lake whitefish, black bears, and gray wolves (*Canis lupus*) also demonstrate the capacity to track spawning migrations of catostomid suckers across landscapes to feed on eggs and spawning adults (Dion & Whoriskey, 1992; Gable et al., 2018; Romain et al., 2013). Thus, spawning migrations of inland fishes like catostomid suckers may produce similar resource dynamics to coastal migratory salmon runs, where inter-specific and inter-population variability in the phenology of spawning fuels a stable supply of eggs and carcasses as resources for mobile consumers (a resource portfolio; Armstrong et al., 2016; Ruff et al., 2011). Interestingly then, regional and latitudinal variation in sucker spawning phenology (Murchie et al., 2024) may also create resource portfolios across whole landscapes that can be exploited by mobile terrestrial and aquatic consumers.

Resource portfolios can increase consumer growth rates and population resilience by allowing consumers to switch between alternative resource species and habitats to maintain growth, even in the face of disturbances or depletion of a single food source or habitat (Hale et al. 2025). The seasonal patterning of spawning events across north-temperate fish species, occurring predominantly in spring and fall, mediates the phenology of egg availability (Figure 5). We propose that phenological diversity in synchronized spawning events facilitates temporally heterogeneous pulses of high-quality resources in inland freshwaters that are predictably patterned through the year. These temporally and spatially explicit resource pulses subsidize microhabitats in which spawning occurs and can be broadly exploited by consumers, creating hotspots of food web activity that mobile consumers can move between to track waves of available egg prey (as seen in terrestrial and coastal mobile predators; Armstrong et al., 2016). Spawning windows are generally viewed as periods of high sensitivity for spawning species (Dahlke et al., 2020), receiving legislated protections in Canada (Tunney et al., 2023). Interestingly though, these windows may also be critical periods of energy flux that shape the growth and survival of egg predators across landscapes.

### 4.4 Comparing reproductive resource pulses and masting events

Masting events have long fascinated the ecological community due to the visible resource hyper-abundance, synchronization, and unpredictability. To suggest implications and new research directions, we explore similarities and differences between aquatic resource pulses in the form of fish eggs and terrestrial resource pulses in the form of seed fall by long-lived plant species. In masting species, large quantities of reproductive material, mast fruit and seeds, are released synchronously to increase offspring survival through targeted predator satiation (i.e., predator swamping), improved reproductive efficiency via pollination, and/or optimal and predictable environmental conditions (Beck et al., 2024). Diverse consumers across trophic guilds (e.g., insects, small mammals, and birds, including those that are rarely seed predators) are attracted to masting events through a “birdfeeder effect”. Fish spawning events also involve the synchronous deposition of reproductive material that swamp predators and may create a similar birdfeeder effect by aggregating diverse and abundant communities of consumers (e.g., egg predators, like fishes, invertebrates, and other aquatic and terrestrial consumers and their predators). However, compared to masting events, spawning may be more predictable for consumers. This improved predictability may facilitate tracking of freshwater fish spawning events across habitats by highly mobile consumers, as consumers exhibit similar behavioural responses to capitalize on resource pulses introduced by both fish spawning and masting (Armstrong et al., 2016; Fuiman et al., 2015; Ostfeld et al., 1996; Ostfeld & Keesing, 2000; Ruff et al., 2011). In both cases, the hyper-abundance of available resources leads to rapid energy flow that propagates through local and regional food webs, with the capacity to alter individual and population growth rates (Ostfeld et al., 1996; Rinella et al., 2012), competitive dynamics (Bailey et al., 2019; Selva et al., 2012), and adaptive foraging by mobile and local predators (Jędrzejewska & Jędrzejewska, 1998; Ruff et al., 2011). Thus, aggregate fish spawning events and inter-annual variation in its intensity may have similar consequences for food webs relative to other more widely studied resource pulses (e.g., masting); the similarity or differences between these pulsed energy flows through food webs deserves future study.

## 5.0 Conclusions

Though spawning events and their exploitation are typically cryptic, here, we combined current molecular approaches, targeted field sampling, and syntheses of extensive natural history observations and published literature to reveal that reproductive resource pulses represent hotspots for food web activity in freshwater systems. In particular, we demonstrated that white sucker spawn is a ubiquitous resource for local and mobile consumers, attracting terrestrial and aquatic invertebrates, fish, mammals, and birds. White sucker are among the most widespread migratory freshwater fish species in North America, suggesting that the trophic hotspots they catalyze are also widespread across inland freshwaters. However, both the provisioning and exploitation of eggs are widespread across freshwater fishes in north-temperate systems; many fish species may provision similar trophic hotspots during their annual bouts of synchronized reproduction. Similar to terrestrial masting events and marine mass-reproductive events, reproductive resource pulses in freshwater systems appear to concentrate consumers in space and time, likely leading to cascading impacts on their broader food webs. Thus, reproductive resource pulses and the hotspots they catalyze are broadly underappreciated but likely critically important for the ecological functioning of freshwater ecosystems, from local food web to landscape processes.

## 6.0 Methods

### 6.1 Field Study

#### 6.1.1 Study Site

Smoke Lake is a long-term study site in Algonquin Provincial Park, Ontario, Canada (45° 30’ 58.28” N, 78° 40’ 55.69” W). The lake has a maximum depth of 55 m, mean depth of 16.2 m, and surface area of 661 ha, with several deltas formed from inflowing creeks. In previous years of sampling the aquatic community, the authors and other staff from the Harkness Laboratory of Fisheries Research observed the presence of suckers at two creek deltas in the early spring, likely representing spawning runs. Additionally, anglers and technicians noted large numbers of smallmouth bass alongside migrating suckers. To quantify the exploitation of sucker eggs in this lake system, we selected three sampling sites located in creek deltas: one site where both migrating white suckers and eggs had been previously observed, one site where only migrating white suckers had been previously observed (upstream movement blocked by a large beaver dam), and one negative control site where no aggregations of adult white suckers or eggs had been observed.

#### 6.1.2 Fish and Invertebrate Sampling

At each of the sites, we sampled multiple levels of the aquatic food web daily for approximately one week after spawning aggregations of white suckers were visible (May 10, 2023 to May 16, 2023). To capture predatory and prey fish species, we used rod-and-reel angling and fish traps (set for 12-16 hours) baited with 50-ml of chicken-flavoured dog kibble within 200 metres of creek deltas. Upon capture, fish were euthanized via cervical dislocation, followed by pithing of the brain. Carcasses were placed into labelled sample bags and kept on ice until full dissection. Within 6 hours of euthanasia, fish were transported to the Harkness Laboratory of Fisheries Research where they were either dissected using aseptic techniques to remove the gastrointestinal tract or frozen whole at -20°C. When dissected, we identified all stomach contents to the finest possible taxonomic resolution. We then recorded the number and batch weight of each diet category, collecting stomach and intestinal contents separately into labelled conical centrifuge tubes (2 mL, 15 mL, or 50 mL; ThermoFisher Scientific, Mississauga, Ontario). All dissected samples and fish that were frozen whole were sent to the Canadian Department of Fisheries and Oceans, Fisheries and Ecosystem Science Division, Gulf Fisheries Centre in Moncton, New Brunswick, Canada, where whole fish were fully dissected and processed as described above and gastrointestinal contents were analyzed using molecular techniques described below. After ∼20 smallmouth bass were collected from a site, any additional fish were non-lethally sampled for stomach contents via gastric lavage as a part of a separate long-term monitoring and tagging program. All lavaged contents were collected into labelled sample bags, placed on ice, and identified and weighed within 6 hours of collection. All animal handling protocols were approved by the University of Toronto Animal Care Committee and performed under the following Animal Use Protocol: #20012677.

Macroinvertebrates were collected from the benthos of each site by sweeping a D-net along the sediment and around overturned cobbles and boulders in nearshore areas (<1 m depth) within 50 metres of creek inflows. To supplement macroinvertebrate samples, we also snorkelled the shoreline within the same radius of the inflow, flipping rocks and inspecting coarse woody debris and macrophytes by hand. We focused on Trichoptera, Plecoptera, Odonata, Ephemeroptera, Megaloptera, and Decapoda, when available. All macroinvertebrates except Decapoda were then placed into labelled sample bags, placed on ice, and frozen at -20°C within 6-hours of capture. Decapoda were dissected using aseptic techniques and the gastrointestinal tracts including their contents were removed, placed into labelled 2 mL microcentrifuge tubes, and frozen at -20°C. All dissected samples and macroinvertebrates that were frozen whole were sent to the Canadian Department of Fisheries and Oceans, Fisheries and Ecosystem Science Division, Gulf Fisheries Centre in Moncton, New Brunswick, Canada, where individuals were stored at -80C until they were thawed on ice immediately before dissection. In the lab, diet contents of both fish and invertebrates were transferred directly into microcentrifuge tubes containing T1 lysis buffer (Macherey-Nagel).

#### 6.1.3 Molecular Analysis of Gastrointestinal Contents Extracted from Fish and Invertebrates

Whole genomic DNA was extracted from the samples using the Macherey-Nagel nucleospin II kit (Macherey-Nagel) with a few minor modifications to the manufacturer’s instructions. These modifications included a brief vortexing every 30-45 minutes during a 3-hour incubation, and a brief spin of the lysate at 6000 x g to pellet undigested debris before transferring to a new tube containing buffer B3.

To design and test the white sucker qPCR assay, a total of 472 COI nucleotide sequences of Catostomus spp., including 86 individuals of white sucker and 77 individuals of the sympatric longnose sucker, were downloaded from GenBank and aligned in Geneious using the MAFFT algorithm. To minimize the potential for off-target amplification, the forward and reverse primers for white sucker were designed to have 4 mismatches with the non-target longnose sucker and the fluorescent probe contains 5 mismatches between the species. Longnose sucker DNA was not available for testing non-specific amplification. PCR reactions were conducted in 20 µl volumes with 1X SsoAdvanced Universal Probes Supermix (Bio-Rad Laboratories,Inc) and 3 µl of genomic DNA with of 600nM of forward primer Ccom_for4(5’-TAAAACCCCCAGCCATCTCTCAA-3’) and 600nM of reverse primer Ccom_rev4(5’-AAATTTCGGTCTGTTAGTAGCATAG-3’) and 250nM of the 5’ fluorescein-labelled minor-groove binding probe Ccom4_COI(56-FAM/CTGTTCTCCTCCTCTTATCATTACCTGT/3MGB-NFQ) from Integrated DNA Technologies. Cycling conditions consisted of an initial denaturation at 95 °C for 3 minutes followed by 45 cycles of 95 °C for 15 seconds and 63 °C for 30 seconds and fluorescence detection.

#### 6.1.4 Fish Telemetry

To investigate whether a mobile consumer tracked the white sucker reproductive resource pulse in space and time, we leveraged fish telemetry data from an existing array located in Smoke Lake (Algonquin Provincial Park, ON, Canada). Acoustic detections from three Vemco VR2W omnidirectional receivers (INNOVASEA Ltd., Bedford, NS, Canada) closest to the sampling sites were selected to investigate species-specific site occupancy. Throughout the study duration, the tagged populations of white sucker and smallmouth bass in Smoke Lake were comprised of 15 and 19 individuals, respectively, carrying uniquely coded 69kHz acoustic telemetry transmitters (INNOVASEA Ltd., Bedford, NS, Canada). Detection data of deceased fish was excluded from all analyses. To compare the abundance of smallmouth bass and white sucker among each of the three sites, we quantified the number of daily detections for both species during May 2023. To identify temporal and spatial association between smallmouth bass and spawning suckers, we calculated the number of unique tagged individuals detected at Site 3 weekly from winter, prior to the sucker spawning migration (March 2023), to early summer, following sucker spawning (July 2023). In choosing this broad window of time, we can capture the potential pre-arrival, residence, and departure of white sucker and smallmouth bass at this delta. If smallmouth bass spatially and temporally track the sucker spawning migration observed in Smoke Lake, then the number of individual white sucker detected at Site 3 should be positively correlated with the number of individual smallmouth bass detected prior, during, and following sucker spawning.

### 6.2 Natural History Observations

#### 6.2.1 Monitoring exploitation of a white sucker spawning migration by terrestrial consumers in Algonquin Provincial Park, Ontario

Historic observations in Algonquin Park suggested that white sucker spawning aggregations were heavily exploited by resident black bears, which motivated the collection of 8 years of standardized natural history observations at a sucker spawning aggregation site in Algonquin Provincial Park, Ontario, Canada. The observations were initially intended to monitor black bear behaviour at a feeding site rich in energy-dense adult sucker and their eggs at a time of year when protein and fat sources are relatively low. The scope of the project was expanded to include all consumers in the first year of the project (2008). From 2008-2015, observations were collected from an observation blind and a network of camera traps at the mouth of Wright Creek in Opeongo Lake, beginning when creek temperatures approached 11°C. Prior to creek temperatures reaching 11°C, all cameras and the observation blind were in place to record activity at the spawning site.

The observation blind was a two-person deer stand enclosed in camouflage canvas placed approximately 5.5 m above the water surface, downstream of the spawning site. The blind was enclosed except for a 30 cm x 120 cm viewing slit, covered in camouflage mosquito screen. We also monitored the stream using a 5-camera system that included Reconyx RapidFire RM45, Uway model VH-GSMB, and HCO ScoutGuard model SG550 to determine species absent during blind observations. All cameras were motion-activated, infrared capable, and oriented to minimize false captures instigated by sunrise and sunset. They were spaced such that they provided as close to complete coverage of the spawning site as possible. All photographs were reviewed by Ontario Ministry of Natural Resources and Forestry staff to identify species present and qualitatively describe observed behaviour.

Blind observations were recorded during three periods of the day: morning (∼7:00 to 10:00), afternoon (∼11:00 to 14:00), and evening (∼17:30 to 20:30). Observation events were scheduled randomly within those windows and predominantly conducted by individual observers. During blind observations, all avian and terrestrial species that could be observed were recorded, regardless of whether they were observed to feed on suckers and eggs. We identified and counted all individuals present at the stream within the field of view every 10 minutes and used the intervening period to carry out additional focal animal sampling. During focal sampling, we detailed subsurface foraging by aquatic birds. Here, we recorded time spent beneath the water’s surface, which was assumed to be reflective of time spent feeding on sucker eggs. This assumption was verified by the presence of eggs in the bill of surfacing birds or sufficient water clarity for direct observations of underwater consumption. Some waterfowl were also observed feeding on associated invertebrates drawn to the eggs and fish carcasses/offal on shore.

#### 6.2.2 Literature Review

To assess the extent that reproductive resource pulses potentially contribute to local food webs, we conducted a literature review and synthesis focusing on documented egg predation among northern freshwater fishes. We queried Google Scholar, Clarivate Web of Science, and Elsevier Scopus with the following search strings: (1) “(“egg predator” OR “egg predation” OR “egg consumption” OR “egg consumer”) AND freshwater AND fish” [Google Scholar], (2) “(“egg pred*” OR “egg con*”) AND freshwater AND fish” [Web of Science, Scopus]. These searches returned a total of 5400, 1663, and 49 documents from Google Scholar, Web of Science, and Scopus, respectively. We scanned the titles of the first 2000 returns from Google Scholar and all of the articles from Web of Science and Scopus. If titles were deemed relevant, then whole articles were searched for the terms “egg” and “egg pred” to locate documentation of observed egg predation events. Reference lists of all relevant articles were also scanned to find additional relevant works. Only egg predation events documented in studies using wild freshwater fishes from north-temperate (> 30°N) freshwater systems were included. We extracted and recorded all egg predation interactions where taxonomic groups were specifically identified for both the egg predator and prey taxa. If the species of eggs consumed eggs were unidentified, observations were not recorded.

To estimate the seasonal availability of fish eggs to potential egg predators, we extracted spawning dates and embryo development times for species in our literature review from the classic reference volume by Scott & Crossman (1998). Because spawning dates and embryo development times vary across latitudes and populations, for this approximation, we used the mid-points of the reported ranges for spawning date and embryo development time. In this synthesis, we only included species that are known to coexist in the Great Lakes drainage basin and provision eggs according to our review; this allows us to approximate a possible “resource portfolio” of egg availability to consumers (i.e., aggregate variance in egg availability reduced by asynchrony in spawning phenologies and embryo development times across species).

### 6.3 Statistical Analysis

To summarize the relative quantity of white sucker DNA present in the gastrointestinal tracts of consumers, we converted each value to a proportion of the average sucker DNA quantity detected from white sucker egg samples taken across all sites. As such, each proportion estimates the extent to which a consumer’s gastrointestinal contents were comprised of white sucker eggs; higher values represent more recent, more direct, or a higher proportion of egg material in the sampled diet. Then, we used cross-correlation to evaluate the degree of correlation and synchrony between detections of smallmouth bass and white sucker at the site of confirmed sucker spawning (Site 3). Cross-correlations computed Pearson correlation coefficients across all possible time lags for weekly detections of individual white sucker and smallmouth bass at the Site 3 receiver. The number of unique individuals detected per week from March to July, 2023 was transformed into a percentage of the total tagged population for each species per week prior to being used in cross-correlations. Finally, we used non-parametric Kruskal-Wallis ANOVA to compare the number of white sucker and smallmouth bass detections at acoustic receivers across sampling sites in Smoke Lake, Ontario, Canada. We then conducted multiple pairwise comparisons using Dunn’s test with Bonferroni p-value adjustment.

## Acknowledgements

The authors would like to thank Darren Smith, Alyssa Andersen, and Lauren Paulic for providing support in the field. The authors would also like to thank Chloé Melanson for support in the laboratory. BCM was supported by an NSERC Alliance, Grant No. ALLRP 556300-20. NSERC CGS and PDF to TJF. NSF Postdoctoral Research Fellowships in Biology Program under Grant No. 2410512 to KRSH. NSF had no role in this project’s development, design and delivery.

## Author contributions

TJF, TDT, and BCM conceived the study, TJF designed the field data collection, TJF, BMS, and SP completed field data collection and assisted with acoustic telemetry logistics, RS processed and analyzed all laboratory samples, DP and MEO designed and completed all natural history observations, BMS and TM curated and analyzed all acoustic telemetry data, TJF conducted the literature review, TJF, KRSH, BMS, and TM conducted all analyses and created all figures (except Figure 4), TJF and KRSH wrote the initial manuscript draft, BMS, RS, SP, JBA, KSM, MEO, DP, MR, BJS, TDT, and BCM provided critical revisions to the manuscript.

## Competing interests

The authors report no competing interests.

## Data availability

All data and code used in the above analyses and to create all non-illustrative figures are available at the following Zenodo repository: https://doi.org/10.5281/zenodo.17234062

## Notes

https://doi.org/10.5281/zenodo.17234062

### Competing Interest Statement

The authors have declared no competing interest.

