## Supporting Information for "Fish spawning events stimulate trophic hotspots across freshwater food webs"

**Tables**

***Table S1.*** *Taxa observed visually (excluding auditory identifications) during blind observations from Wright Creek, Opeongo Lake, Algonquin Provincial Park, Ontario Canada. Observations were made during late spring (late April to late May) and repeated annually from 2010 to 2015.*

| **Taxa** | **Species** | **2010** | **2011** | **2012** | **2013** | **2014** | **2015** | **Total** | **Percentage of Total Observations** |
| --- | --- | --- | --- | --- | --- | --- | --- | --- | --- |
| **Bird** | Common Merganser | 7 | 61 | 1 | 18 | 55 | 119 | 261 | 38.0 |
|  | Herring Gull | 59 | 5 |  |  |  | 34 | 98 | 14.3 |
|  | American Black Duck |  |  |  | 8 | 26 | 54 | 88 | 12.8 |
|  | Belted Kingfisher | 3 | 4 | 20 | 1 | 7 | 15 | 50 | 7.3 |
|  | Common Raven | 4 | 1 | 18 | 3 | 4 | 9 | 39 | 5.7 |
|  | Bald Eagle | 4 | 1 | 13 |  | 5 | 2 | 25 | 3.6 |
|  | American Crow | 4 |  |  |  | 1 | 20 | 25 | 3.6 |
|  | Turkey Vulture | 9 |  |  | 3 |  | 7 | 19 | 2.8 |
|  | Mallard | 1 |  |  | 17 |  |  | 18 | 2.6 |
|  | Wood Duck | 5 |  | 6 | 3 | 1 |  | 15 | 2.2 |
|  | Gull |  |  | 1 | 3 | 3 |  | 7 | 1.0 |
|  | Spotted Sandpiper | 4 |  |  |  | 3 |  | 7 | 1.0 |
|  | Blue-Winged Teal |  |  |  | 3 | 1 |  | 4 | 0.6 |
|  | Blue Jay |  | 1 |  |  | 1 | 1 | 3 | 0.4 |
|  | Common Grackle | 1 | 1 |  |  |  |  | 2 | 0.3 |
|  | Corvids |  |  |  |  |  | 2 | 2 | 0.3 |
|  | Osprey |  |  | 2 |  |  |  | 2 | 0.3 |
|  | Sandpiper | 2 |  |  |  |  |  | 2 | 0.3 |
|  | Ruby-Throated Hummingbird |  |  |  |  |  | 1 | 1 | 0.1 |
|  | Ring-billed Gull |  | 1 |  |  |  |  | 1 | 0.1 |
|  | Yellow-bellied Sapsucker |  |  |  |  |  | 1 | 1 | 0.1 |
|  | Sparrow |  |  |  |  | 1 |  | 1 | 0.1 |
|  | Winter Wren |  |  |  |  |  | 1 | 1 | 0.1 |
| **Mammal** | River Otter |  | 3 |  |  |  | 2 | 5 | 0.7 |
|  | Raccoon |  |  |  | 4 |  |  | 4 | 0.6 |
|  | Marten |  |  |  |  | 1 | 2 | 3 | 0.4 |
|  | Muskrat |  |  | 2 |  |  |  | 2 | 0.3 |
|  | Red Squirrel |  |  |  |  | 1 |  | 1 | 0.1 |

***Table S2.*** *Taxa observed during camera trap surveys from Wright Creek, Opeongo Lake, Algonquin Provincial Park, Ontario Canada. Observations were made from late spring (late April to late May) to approximately mid-June and repeated annually from 2008 to 2015.*

| **Taxa** | **Species** | **2008** | **2009** | **2010** | **2011** | **2012** | **2013** | **2014** | **2015** | **Total** |
| --- | --- | --- | --- | --- | --- | --- | --- | --- | --- | --- |
| **Bird** | Gull | 825 | 505 | 90 | 12 | 291 | 1 |  | 11 | 1735 |
|  | Common Merganser | 125 | 63 | 4 | 13 | 41 | 6 | 4 | 92 | 348 |
|  | Common Raven | 60 | 28 | 29 | 8 | 37 | 16 | 32 | 48 | 258 |
|  | Bald Eagle |  | 31 | 50 | 18 | 28 | 48 | 5 | 43 | 223 |
|  | Turkey Vulture | 32 | 23 | 31 | 5 | 50 | 8 | 4 | 20 | 173 |
|  | Great Blue Heron | 79 | 27 | 1 |  | 2 |  |  | 11 | 120 |
|  | American Crow | 19 | 4 | 7 | 3 | 3 | 4 | 1 | 17 | 58 |
|  | Wood Duck | 6 | 2 |  | 3 | 20 | 1 |  |  | 32 |
|  | American Black Duck | 7 |  |  |  |  | 8 |  | 7 | 22 |
|  | Wild Turkey |  |  |  | 1 |  | 3 |  |  | 4 |
|  | Canada Goose | 2 |  |  |  |  |  |  |  | 2 |
|  | Blue Jay |  |  |  |  |  |  | 1 |  | 1 |
|  | Broad-winged Hawk |  |  | 1 |  |  |  |  |  | 1 |
|  | Mallard | 1 |  |  |  |  |  |  |  | 1 |
|  | Northern Flicker |  |  |  |  |  | 1 |  |  | 1 |
| **Mammal** | Raccoon | 2 | 26 | 11 | 2 | 1 | 56 | 25 |  | 123 |
|  | Black Bear | 19 | 6 | 66 | 4 | 2 | 2 | 5 | 8 | 112 |
|  | White-tailed Deer | 7 | 8 | 3 | 3 | 10 | 9 |  | 14 | 54 |
|  | Fisher |  |  |  |  |  | 1 | 2 | 27 | 30 |
|  | Red Fox | 6 | 13 |  |  |  |  |  |  | 19 |
|  | Marten |  | 1 |  |  |  |  |  | 10 | 11 |
|  | Moose |  |  | 1 | 2 |  |  |  | 5 | 8 |
|  | Canid | 2 |  |  |  |  |  | 2 |  | 4 |
|  | Algonquin Wolf |  |  |  |  |  |  |  | 1 | 1 |
|  | Muskrat |  |  |  |  |  |  |  | 1 | 1 |
|  | River Otter |  |  | 1 |  |  |  |  |  | 1 |

***Table S3.*** *Summary table identifying egg predators and provisioners of egg resources following a review of the literature conducted using both Google Scholar and ISI Web of Science (see the main manuscript for search string details).*

| **Egg Predator** | **Egg Provisioner** | | **Location** | **Ref** | |
| --- | --- | --- | --- | --- | --- |
| Alewife | Alewife | | Lake Michigan | (Edsall, 1964) | |
| Atlantic salmon | Steelhead | | Salmon River, New York | (Johnson et al., 2016) | |
| Bluegill sunfish | Smallmouth bass | | Ontario, Canada | (Gravel & Cooke, 2009) | |
|  | Common carp | | Minnesota, USA | (Silbernagel & Sorensen, 2013) | |
|  | Bluegill sunfish | | Lake Opinicon, Ontario, Canada | (Gross & MacMillan, 1981) | |
| Bonneville whitefish | Bonneville cisco* | | Utah, USA | (Bouwes & Luecke, 1997) | |
| Brook trout | Steelhead | | Salmon River, New York, USA | (Johnson, 1981) | |
| Brown bullhead | Lake trout | | Lake Opeongo | (Martin, 1957) | |
|  | Round whitefish | | Newfound Lake, Bristol, New Hampshire | (Normandeau, 1969) | |
| Brown trout | Lake sturgeon | | Peshtigo River, Wisconsin, USA | (Caroffino et al., 2010) | |
| Burbot | Lake trout | | Lake Ontario | (Stauffer & Wagner, 1979) | |
|  | European whitefish | | Lake Constance | (Rösch & Schmid, 1996) | |
|  | Round whitefish | | Newfound Lake, Bristol, New Hampshire | (Normandeau, 1969) | |
| Channel catfish | Lake whitefish | | Lake Huron | (Kalejs et al., 2022) | |
|  | Walleye | | Lake Huron | (Kalejs et al., 2022) | |
| Common carp | Walleye | | Lake Huron | (Kalejs et al., 2022) | |
|  | Lake whitefish | | Lake Huron | (Kalejs et al., 2022) | |
|  | Bonneville cisco* | | Bear Lake, Utah | (Bouwes & Luecke, 1997) | |
|  | Lake sturgeon | | Peshtigo River, Wisconsin, USA | (Caroffino et al., 2010) | |
| Deepwater sculpin | Bloater | | Lake Michigan | (Mychek‐Londer et al., 2013) | |
|  | Deepwater sculpin | | Lake Michigan | (Mychek‐Londer et al., 2013) | |
| Dusky shiner | Redbreast sunfish | | South Caronlina, USA | (Fletcher, 1993) | |
| Emerald shiner | Alewife | | Lake Michigan | (Edsall, 1964) | |
| European sculpin | Atlantic salmon | | River Vindelalven | (Palm et al., 2009) | |
| Fallfish | Oncorhynchus spp. | | Salmon River, New York, USA | (Johnson et al., 2009) | |
| Fathead minnow | Fathead minnow | | Meanook, Alberta, Canada | (Vandenbos et al., 2006) | |
| Johnny darter | Walleye* | | Lake Erie | (Roseman et al., 1996) | |
| Lake sturgeon | Lake sturgeon | | Peshtigo River, Wisconsin, USA | (Caroffino et al., 2010) | |
| Lake trout | Lake trout | | Lake Ontario | (Stauffer & Wagner, 1979) | |
| Lake whitefish | Cisco | | Lake Superior | (Stockwell et al., 2014) | |
|  | Longnose sucker | | Gouin Reservoir, Quebec | (Dion & Whoriskey, 1992) | |
|  | White sucker | | Gouin Reservoir, Quebec | (Dion & Whoriskey, 1992) | |
|  | Coregonids* | | Lake Superior | (Woodard et al., 2021) | |
| Lepomis spp. | Rock bass | | Lake Opinicon, Ontario | (Gross & Nowell, 1980) | |
| Logperch | Walleye* | | Lake Erie | (Roseman et al., 1996) | |
|  | Lake sturgeon | | St. Lawrence River, Ontario, Canada | (Johnson et al., 2006) | |
| Longnose sucker | Lake trout | | Lake Ontario | (Stauffer & Wagner, 1979) | |
|  | White sucker | | Gouin Reservoir, Quebec | (Dion & Whoriskey, 1992) | |
| Mottled Sculpin | Lake trout | | Lake Ontario | (Stauffer & Wagner, 1979) | |
| Northern hogsucker | Carpsucker | | Savannah River, Georgia | (Grabowski & Isely, 2007) | |
|  | Spotted sucker | | Savannah River, Georgia | (Grabowski & Isely, 2007) | |
|  | Robust redhorse | | Savannah River, Georgia | (Grabowski & Isely, 2007) | |
|  | Notchlip redhorse | | Savannah River, Georgia | (Grabowski & Isely, 2007) | |
|  | Lake sturgeon | | Peshtigo River, Wisconsin, USA | (Caroffino et al., 2010) | |
| Pumpkinseed sunfish | Smallmouth bass | | Ontario, Canada | (Gravel & Cooke, 2009) | |
|  | Centrarchid spp. | | St. Lawrence River, Ontario, Canada | (Leblanc et al., 2020) | |
|  | Pumpkinseed sunfish | | Lake Banyoles, Spain | (García‐Berthou & Moreno‐Amich, 2000) | |
|  | Bluegill sunfish | | Lake Opinicon, Ontario, Canada | (Gross & MacMillan, 1981) | |
| Quillback | Walleye | | Lake Erie | (Roseman et al., 2006) | |
| Rainbow trout | Steelhead | | Salmon River, New York, USA | (Johnson, 1981) | |
| Redhorse sucker | Lake sturgeon | | St. Lawrence River, Ontario, Canada | (Johnson et al., 2006) | |
| Rock bass | Walleye | | Lake Erie | (Roseman et al., 2006) | |
|  | Lake sturgeon | | Peshtigo River, Wisconsin, USA | (Caroffino et al., 2010) | |
|  | Centrarchid spp. | | St. Lawrence River, Ontario, Canada | (Leblanc et al., 2020) | |
| Round goby | Walleye | | Lake Erie | (Roseman et al., 2006) | |
|  | Smallmouth bass | | Lake Erie | (Steinhart et al., 2004) | |
|  | Nase | | River Rhine, Switzerland | (Lutz et al., 2020) | |
|  | Centrarchid spp. | | St. Lawrence River, Ontario, Canada | (Leblanc et al., 2020) | |
| Round whitefish | Lake trout | | Lake Ontario | (Stauffer & Wagner, 1979) | |
|  | Lake trout | | Lake Opeongo | (Martin, 1957) | |
|  | Round whitefish | | Newfound Lake, Bristol, New Hampshire | (Normandeau, 1969) | |
| Ruffe | Lake cisco | | Lake Superior | (Selgeby, 1998) | |
|  | European whitefish | | Lake Constance | (Rösch & Schmid, 1996) | |
|  | Powan | | Loch Lomond | (Adams & Tippett, 1991) | |
|  | Vendace | | Bassenthwaite Lake, UK | (Winfield et al., 1998) | |
| Sculpin spp. | Walleye* | | Lake Erie | (Roseman et al., 1996) | |
| Silver chub | Walleye | | Lake Erie | (Roseman et al., 2006) | |
| Slimy sculpin | Lake trout | | Lake Ontario | (Stauffer & Wagner, 1979) | |
|  | Cottus sp. | | Lake Ontario | (Owens & Dittman, 2003) | |
|  | Bloater | | Lake Michigan | (Mychek‐Londer et al., 2013) | |
|  | Deepwater sculpin | | Lake Michigan | (Mychek‐Londer et al., 2013) | |
| Spottail shiner | Walleye | | Lake Erie | (Roseman et al., 2006; Wolfert et al., 1975) | |
|  | Alewife | | Lake Michigan | (Edsall, 1964) | |
| Stonecat | Walleye | | Lake Erie | (Wolfert et al., 1975) | |
| Three-spine stickleback | Pelagic whitefish | | Lake Constance, Switzerland | (Baer et al., 2021) | |
|  | Northern pike | | Kalmar Sound, Baltic Sea | (Nilsson, 2006) | |
|  | Three-spine stickleback | | Lake Wapato, Washington, USA | (Semler, 1971) | |
| Trout-perch | Walleye* | | Lake Erie | (Roseman et al., 1996) | |
| Utah sucker | Bonneville cisco* | | Bear Lake, Utah | (Bouwes & Luecke, 1997) | |
| Walleye | Walleye | | Lake Huron | (Kalejs et al., 2022) | |
| White perch | Walleye | | Lake Erie | (Roseman et al., 1996; Schaeffer & Margraf, 1987) | |
|  | White bass | | Lake Erie | (Schaeffer & Margraf, 1987) | |
|  | White perch | | Lake Erie | (Schaeffer & Margraf, 1987) | |
| White sucker | Lake trout | | Lake Ontario; Lake 468, ELA | (Stauffer & Wagner, 1979; Wasylenko et al., 2013) | |
|  | Longnose sucker | | Gouin Reservoir, Quebec | (Dion & Whoriskey, 1992) | |
|  | Walleye | | Lake Erie | (Roseman et al., 2006; Wolfert et al., 1975) | |
|  | Lake sturgeon | | Peshtigo River, Wisconsin, USA | (Caroffino et al., 2010) | |
| Yellow perch | Round whitefish | | Newfound Lake, Bristol, New Hampshire | (Normandeau, 1969) | |
|  | Lake whitefish | Lake Huron | | | (Kalejs et al., 2022) |
|  | Walleye | Lake Huron, Lake Erie | | | (Kalejs et al., 2022; Roseman et al., 1996) |
|  | Lake trout | Keuka Lake, New York, USA | | | (Fitzsimons, 1990) |
|  | Lake sturgeon | Peshtigo River, Wisconsin, USA | | | (Caroffino et al., 2010) |
|  | Centrarchid spp. | St. Lawrence River, Ontario, Canada | | | (Leblanc et al., 2020) |
